# The Q1K Integrated EEG and Eye-Tracking Experimental Test Battery for Open Autism Science

**DOI:** 10.64898/2026.07.13.738272

**Authors:** Christian O’Reilly, James Desjardins, Gabriel Blanco-Gomez, Scott Huberty, Anthony Hosein Poitras Loewen, Sweety Ramnani, Diksha Srishyla, Inga S Knoth, Charles-Olivier Martin, Jonathan P Batten, Stefon van Noordt, Noemie Hebert-Lalonde, Fabienne Samson, Tim J Smith, Baudouin Forgeot d’Arc, Carl Ernst, Melissa T Carter, Christine Lucas Tardif, Julie Scorah, Mélanie Couture, Pascale Abadie, Ridha Joober, Sébastien Jacquemont, Guy A Rouleau, Sarah Lippe, Mayada Elsabbagh

**Affiliations:** Computer Science and Engineering, University of South Carolina, Columbia, SC, USA; Artificial Intelligence Institute, University of South Carolina, Columbia, SC, USA; Carolina Autism and Neurodevelopment Research Center, University of South Carolina, Columbia, SC, USA; Institute for Mind and Brain, University of South Carolina, Columbia, SC, USA; Montreal Neurological Institute-Hospital, Montreal, Canada; Compute Ontario, St. Catharines, Canada; Department of Psychology, University of Montreal, Montreal, QC, Canada; CHU Sainte-Justine Research Centre, University of Montreal, Montreal, QC, Canada; SR Research, Ottawa, Canada; Department of Psychology, Wake Forest University, Winston-Salem, NC, USA; Creative Computing Institute (CCI), University of the Arts London, UK; Centre of Brain and Cognitive Development (CBCD), Birkbeck University of London, UK; Département de Psychiatrie et d’addictologie, Université de Montréal, Montreal, QC, Canada; Centre de recherche Azrieli du CHU Sainte-Justine, Montreal, QC, Canada; Department of Human Genetics, McGill University, Montreal, QC, Canada; Department of Genetics, Children’s Hospital of Eastern Ontario, University of Ottawa, Ottawa, ON, Canada; Department of Neurology & Neurosurgery, McGill University, Montreal, Quebec, Canada; Department of Biomedical Engineering, McGill University, Montreal, Quebec, Canada; Occupational Therapy Program, School of Rehabilitation, Université de Sherbrooke, Sherbrooke, Canada; Child and Adolescents Psychiatry Division, Department of Psychiatry, Rivière-des-Prairies Mental Health Hospital, CIUSSS-NIM, Montreal, Quebec, Canada; Douglas Mental Health University Institute, Montreal, Quebec, Canada; Department of Psychiatry, McGill University, Montreal, Quebec, Canada; Department of Pediatrics, University of Montreal, Montreal, QC, Canada

**Keywords:** Autism spectrum disorder, open-science, eye tracking, electroencephalogram, multimodality

## Abstract

The Quebec 1000 Families (Q1K) platform has been designed to recruit, phenotype, and collect biospecimens from a large cohort of families with at least one member with autism spectrum disorder or a related neurodevelopmental condition. As part of the Q1K protocol, an experimental test battery was developed and validated using simultaneous high-density electroencephalography (EEG) and eye tracking (ET). We report on the general approach and design principles, multimodal EEG/ET integration, the tasks, and their validation, providing a blueprint for implement such a project. It also describes the cohort and its methods as a reference for future studies using this dataset. By releasing openly this experimental test battery, we aim to support task standardization and multi-project data pooling in autism and related neurodevelopmental disorders.

## 1. Introduction

Although the issue of non-replicability of research results is general in science (Aarts et al., 2015; Baker, 2016) and has often been driven by insufficient sample sizes (Szucs & Ioannidis, 2017), it is particularly challenging for fields like autism where research samples are often characterized by high heterogeneity. Indeed, participant heterogeneity is a fundamental property of autism, a condition defined as a *spectrum* explicitly to include a range of different etiologies and symptom profiles. When this substantial etiological and phenotypic heterogeneity is not explicitly modeled, latent subgroup structure and unmeasured moderators can interact with measured variables, producing confounding effects and contributing to inconsistent and poorly replicable findings across studies (Happé et al., 2006; Lenroot & Yeung, 2013; Yarkoni, 2022). Approaches to address these confounders, such as stratification, require larger sample sizes. To address this need, multiple projects involving large cohorts of autistic participants have been conducted over the years. The Autism Brain Imaging Data Exchange (ABIDE) I (Di Martino et al., 2014) and II (Di Martino et al., 2017) are good examples of large neuroimaging cohorts in autism, containing together over 2000 participants scanned with Magnetic Resonance Imaging (MRI). The Infant Brain Imaging Study (IBIS) also collected a large sample of MRI data as well as electroencephalogram (EEG) to provide a complementary assessment of neural activity in autism (Dickinson et al., 2024). Other projects, like the EU-AIMS Longitudinal European Autism Project (LEAP), focused more on a clinical characterization (including MRI) of a large (N>700) full-spectrum (IQ from 50 to 148) sample of participants, or focused on genetic data, like the Autism Speaks MSSNG Project (over 5000 genetic samples from families with an autistic member) (Yuen et al., 2017).

Although these projects have produced a wealth of information on autism, datasets providing a deep phenotyping covering a comprehensive set of modalities (e.g., neuroimaging, behavioral, clinical, and genetic) and inclusive of people across the autism spectrum, along with their relatives, are rare. Quebec 1000 Families (Q1K) was designed to fill this gap. Our aim is to introduce the open science resources it developed and released, with a focus on its integrated EEG and eye tracking (ET) battery. Designed to capture the full spectrum of autism and related conditions, Q1K includes participants of all ages from five years of age onward, ranging from individuals with average cognitive ability and mild support needs to those with more profound symptoms and high support needs, as well as their family members. From its inception, Q1K has been envisioned as an open science initiative, committed to providing a rich, openly accessible dataset. This resource is intended to support the research community in exploring a wide range of questions related to autism and other neurodevelopmental disorders.

Q1K comprises three phases involving increasing levels of phenotyping, with the possibility for the participating families to be included or not in subsequent phases. The first phase involved the creation of a large (target sample: 1,000 families) Registry of people from the province of Quebec, Canada, interested in participating in research projects related to autism. This phase collected relatively shallow information, including demographics and consent to be recontacted. The second phase included light phenotyping, involving mainly the collection of biosamples for genetic analyses as well as questionnaire data. Finally, the third phase (i.e., the deep phenotyping phase) was designed to collect comprehensive multimodal data, including functional and structural 7T MRI, simultaneous EEG and ET (EEG/ET), and comprehensive behavioral characterization. This paper focuses on the experimental test battery developed for the EEG/ET part of the project. It aims to share its conceptual design and describe its EEG/ET-related tools and data to guide researchers in planning to use these resources.

## 2. Methods

### 2.1. Rationale for task selection

A series of consultations with various research teams led to the selection of measures that would best advance understanding of neural mechanisms in autism. Several studies suggest altered sensory processes, brain rhythm entrainment, stimulus discrimination, and brain connectivity in autism and autism-related populations (Arutiunian et al., 2024; Hudac et al., 2018; Lalancette et al., 2023; O’Reilly et al., 2017; Proteau-Lemieux et al., 2025; Samoylov et al., 2024; Shou et al., 2017). Yet, we still need to disentangle the direction of the associations between genotypes, brain dynamics, and behavioral phenotypes in autistic populations (i.e., Duffy & Als, 2019). To design a test battery that addresses this challenge, a few guiding principles were considered in selecting the tasks and designing the battery.

#### Wide coverage

The proposed EEG Q1K protocol includes structured tasks (e.g., oddball paradigm) and naturalistic or non-task conditions that assess autonomic control, sensory processing, neuronal entrainment, discrimination, and attention. Conditions have been designed to cover various experimental paradigms, including naturalistic, resting state, steady-state entrainment, and event-related responses. In turn, these paradigms allow for a wide range of analyses, including evoked potentials, spectral, and time-frequency analyses of brain rhythms, as well as connectivity analyses. Further, the availability of 7T MRI data will provide the opportunity for high-resolution head models, allowing precise EEG cortical and subcortical source estimation as well as multimodal analyses (e.g., integrating or comparing structural and functional connectivity and correlating it with genetics).

#### Established tasks

Conditions and task design were aligned with other cohorts of autism-related conditions, e.g., EU-AIMS LEAP (Loth et al., 2017). Contrary to most typical research projects, the objective of Q1K was not to answer specific research hypotheses. Therefore, in general, task selection and implementation did not involve finessing existing experimental designs or creating novel tasks to address such hypotheses. Rather, the Q1K philosophy is aligned with producing a rich multimodal dataset that can be used openly to support data-driven analyses and answer a wide range of questions on autism. It aims to produce a deep phenotypical characterization of a large number of participants using well-established tasks. The project also decided to reuse well-established tasks to facilitate data pooling with other cohorts, supporting analyses requiring large samples (e.g., studying the heterogeneity of this population and establishing clinically meaningful autism subtypes using machine learning).

#### Whole-spectrum inclusion

Relatively simple tasks not requiring instructions or language skills were prioritized to ensure that participants of almost all ages (5-89 years of age^1^) and across the spectrum can perform these tasks. Thus, we selected tasks developed and tested on toddlers and children. No instructions were provided for any task to minimize the dependence on language understanding and compliance with instructions.

#### Multimodality

This experimental battery was designed for simultaneous and integrated EEG and ET. It includes classical EEG tasks (e.g., Visual and Auditory Steady-State tasks) and classical ET tasks (e.g., Gap Overlap) and provides both modalities for most of these tasks. More specifically, EEG was used for all tasks, while ET was not recorded for some tasks where it was less relevant and was complicating the protocol due to the need for recalibration of the ET system.

### 2.2 Task ordering

EEG/ET recordings are conducted over a one-hour session. Although, in the absence of technical complications, it is possible to complete this protocol within an hour with compliant participants and experienced experimenters, we did not expect to be able to run the whole protocol with all participants, for example, with young children or individuals with more severe symptoms profiles. Thus, we ordered the protocol to run “core” tasks (tasks with the highest priority) at the beginning and “optional” tasks (e.g., those that provided less convincing evidence during piloting) toward the end of the protocol. When ordering tasks, we also considered the comfort of the participants by avoiding running back-to-back boring (e.g., Tone Oddball), passive (e.g., Resting-State), or “abrasive” tasks (e.g., Pupillary Light Reflex and the Visual Steady-State tasks). We ran the resting state first to avoid “spillover” effects from more engaging/stimulating tasks on oscillatory measures. The task order is shown in Table 1.

**Table 1.** The Q1K EEG/ET Task Battery: task order, associated construct, related primary reference, structure (trials and blocks), and approximate duration.

| # | Task | Construct(s) | Primary reference | Trials X blocs | Duration (minutes) |
| --- | --- | --- | --- | --- | --- |
| 1 | Resting-State (RS) | *General brain state* | (Neuhaus et al., 2021)7/13/26 3:10:00 PM | 1-minute X 4 | 5 |
| 2 | Tone Oddball (TO) | *Sensory/Language/Attention* | (Green et al., 2020) | 60 X 1 (450 tones) | 8 |
| Eye tracking calibration |  |  |  |  |  |
| 3 | Gap Overlap (GO) | *Attention/executive function* | (Portugal et al., 2021) | 36 X 1 | 4 |
| Impedance check-up |  |  |  |  |  |
| 4 | Visual Steady-State Response (VSSR) | *Sensory/perception* | (Lalancette et al., 2022) | 40 X 2 | 5 |
| 5 | Auditory Steady-State Response (ASSR) | *Sensory/perception* | (Edgar et al., 2016) | 20 X 1 | 7 |
| Impedance check-up |  |  |  |  |  |
| 6 | Naturalistic Social Preference (NSP) | *Social motivation* | (Saez de Urabain et al., 2017) | 10 videos | 4 |
| 7 | Pupillary Light Reflexes (PLR) | *Sensorimotor* | (Nyström et al., 2018) | 12 X 1 | 2 |
| 8 | Visual Search (VS) | *Perception* | (Gliga et al., 2015) | 30 X 1 | 3 |

### 2.3 Tasks

Below, we describe the tasks included in the battery and lay out the selection rationale. Since we aimed to reuse established tasks to facilitate future between-project data pooling, we selected a *primary reference* for each task (see Table 1). We replicated the protocol from these primary references as closely as possible. Whenever appropriate, we used the results from these reference studies to validate our task implementation during the pilot phase. Supplementary Table 2 lists events common across tasks, whereas Supplementary Tables 3-10 list events specific to each task.

#### Resting-state

Resting-state recordings provide a baseline for the other tasks. These recordings can also reveal various features of cortical network dynamics (e.g., processes associated with the default mode network, power spectral density profiles, cross-region/frequency synchronization) potentially relevant to autism and neurodevelopmental disorders. It also provides an easier paradigm to pool across experiments for future re-use in meta-analyses, metabases, and similar contexts.

For this task, we used an eye-open paradigm to reduce artifacts from eye movements naturally occurring in the eye-closed condition. We followed the protocol established in the GENDARR study (Neuhaus et al., 2021). It consisted of four different abstract videos that looked like screen savers and played in the center of the participant’s screen. Such abstract videos are commonly used in resting-state experiments with children to help keep the participants steady and quiet. To minimize eye movements, videos were relatively small, covering approximately 9.1° (width) × 6.2° (height) of the visual field. Each video lasted 1 minute, with an opportunity for a break (experimenter-controlled, normally about 20 s) between each video. Up to two extra blocks with additional videos could be run if the experimenter judged that the data contained too many noisy periods and required longer recording to obtain sufficient clean data. Videos were selected in random order.

#### Tone oddball

The oddball paradigm involves an event-related task using frequent (standard) and infrequent (odd, deviant) stimuli. It is typically used to study how the brain reacts to unexpected stimuli, for example, by looking at event-related potentials (ERP) such as the mismatch negativity (MMN) and the P150 component of auditory sensory processing. Multiple studies have used the MMN to assess abnormal auditory processing (Green et al., 2020). Both differences in amplitude and latency of the MMN component have been shown in autistic children and adults (Chen et al., 2020; Green et al., 2020; Schwartz et al., 2018). Overall, these studies have shown that failure to correctly perceive sounds (tones) and speech (vowels, syllables) strongly correlates with language deficits (Seery et al., 2013).

Language deficits are common in autistic children, with the proportion of minimally verbal children varying between 25% and 35%, depending on the reports (Rose et al., 2016). The mechanisms behind language deficits are yet to be fully understood but are believed to involve a combination of differences in brain connectivity, difficulties acquiring social abilities, and atypical auditory processing (Matsuzaki et al., 2019). It is widely believed that disruptions of the basic auditory processing processes (feature discrimination, source identification, phonological memory, etc.) are highly tied to general language deficits in autism and other common language disorders (Roberts et al., 2011). Therefore, this task was included in the Q1K battery given the prevalence of language impairments in autism and the language correlates of auditory oddball with low-level mechanisms underlying language.

For this task, two sinusoidal tones (standard [26/30; 86.6%]: 1000 Hz; deviant [4/30; 13.3%]: 2000 Hz) lasting 70 ms (10 ms waxing, 50 ms plateau, 10 ms waning) were played with a 1000 ms interstimulus interval (ISI). Trials were presented in a pseudorandom order. During the task, the Disney short film *Bao* was played without sound. We originally piloted this task using the same abstract videos used for the Resting-State task, but participants found it too difficult to maintain focus. We used one block lasting 8 minutes and containing 60 trials. Each trial consists of 4 to 9 standard tones followed by a deviant tone, for a total of 450 tones across the 60 trials. Each trial is separated by 1000 ms (i.e., the same duration as the ISI).

#### Gap Overlap

The Gap Overlap task is a classical ET task. It starts with a central fixation target animation (a rotating clock). A peripheral stimulus (a rotating cloud) is then presented at the same time (baseline), before (overlap), or after (gap) the disappearance of the central fixation target. A reward animation (a waxing and waning star) is then presented at the same place as the target animation when the participant gaze at the peripheral stimulus. This task is designed to characterize attention to “stickiness”. Evidence suggests that atypical development of visual attention early in life is associated with later developmental outcomes, including autism (Elsabbagh et al., 2013). These atypicalities have been linked to the development of the oculomotor and attention systems (Saez de Urabain et al., 2017). The Gap Overlap task offers a way to measure flexibility in attention switching by quantifying the cost of disengaging from a central stimulus to focus on a peripheral one, which was reported to be impaired in autistic children and adults (Kawakubo et al., 2007; Landry & Bryson, 2004).

In our protocol, the gap conditions consisted of a peripheral target that started 200 ms after the central fixation animation disappeared from the screen. In the overlap condition, the fixation animation stopped rotating at the time of the peripheral target onset, and the fixed image central fixation stayed on the screen until the gaze reached the target region of interest. Finally, in the baseline condition, the peripheral stimulus appeared immediately after the central fixation disappeared from the screen. A single block was used, with 36 trials in the following factorial design: 3 conditions (baseline, gap, overlap) X 2 sides (peripheral stimulus on the left/right side of the central fixation) X 6 repetitions. Trials were recycled if they were invalid because the participant did not look at the fixation stimulus for 500 ms (distracted), took more than 1200 ms to look at the peripheral target (distracted), had a saccade reaction time of less than 100 ms (preempted the apparition of the peripheral target), or gaze on the wrong side of the screen. Whenever the participants failed to gaze at the peripheral target within 2,500 ms, the reward stimulus was displayed as though a gaze at the target had been detected, and the task proceeded to the next trial.

#### Visual Steady-State Response

Differences in sensory processing and perceptual functions have been identified in various neurodevelopmental disorders and are recognized as an important feature of autism (Simon & Wallace, 2016). Typically, these differences are characterized by measuring neural synchrony (Lajiness-O’Neill et al., 2018). Findings from previous research are mixed regarding what frequency bands are relevant and whether autistic individuals differ on baseline and baseline-correct evoked responses (e.g., Orekhova et al., 2007; Port et al., 2016).

An alternative approach is to present oscillatory stimuli at defined frequencies of interest. This method allows directly examining the resonant properties of stimulus processing in the sensory cortex and increasing the signal-to-noise ratio at specific frequencies. We included two tasks to study resonant properties in the visual (VSSR) and auditory (ASSR) cortex.

Disruptions in the rhythmic synchronization of neural activity to visual stimuli could contribute to alterations in sensory processing in autism. Previous research has shown atypical visual steady-state responses in autism in several frequency bands (Simon & Wallace, 2016) and increased short-range lateral inhibition with greater autistic symptom severity (Dickinson et al., 2018). Disruption in the temporal coordination of low-level visual responses could implicate large-scale network integration, which is thought to be compromised in autism (Uhlhaas & Singer, 2012). Examining basic visual steady-state responses also provides valuable data for understanding cortical entrainment and specifically for processing more complex visual stimuli (e.g., faces).

For this task, we used 700 ms videos of circular random sand patterns alternating between 180° rotated copies at rates of 6 Hz or 15 Hz. Multiple combinations of contrast (50% vs 100% of the white-to-black range), spatial resolution (i.e., grain size; 200 vs 400 grain diameter), presentation rates (6 Hz, 10 Hz, and 15 Hz), and the mode of pattern alternation (180° rotations vs luminance inversion) were piloted (Figure 1). We found that the most salient neural responses were obtained for low spatial resolution (i.e., low spatial frequencies in the sand pattern), high contrast, and luminance inversion, which unfortunately corresponded to the conditions that were the least comfortable for the participant (unpublished results). We adopted the experimental conditions that provided the best tradeoff between being not too abrasive and providing high inter-trial coherency^2^. A spatial resolution of 200 dots (i.e., sand grains) across the circle diameter was selected with 100% contrast (range from back to white). The presentation rates were selected to maximize the between-condition contrast in the EEG spectral responses at fundamental and harmonic frequencies. Two blocks are presented, with 40 trials per block. Each trial has a 500 ms pause with a red central fixation ‘+’ followed by a 500 ms central attention animation (a star zooming in and out), a jittered 1100-1500 ms pause displaying the fixation screen, and 700 ms of alternating stimulus presentation. The experiment allows for the experimenters to run an extra block if they consider that one of the blocks is invalid or unreliable.

**Figure 1.**
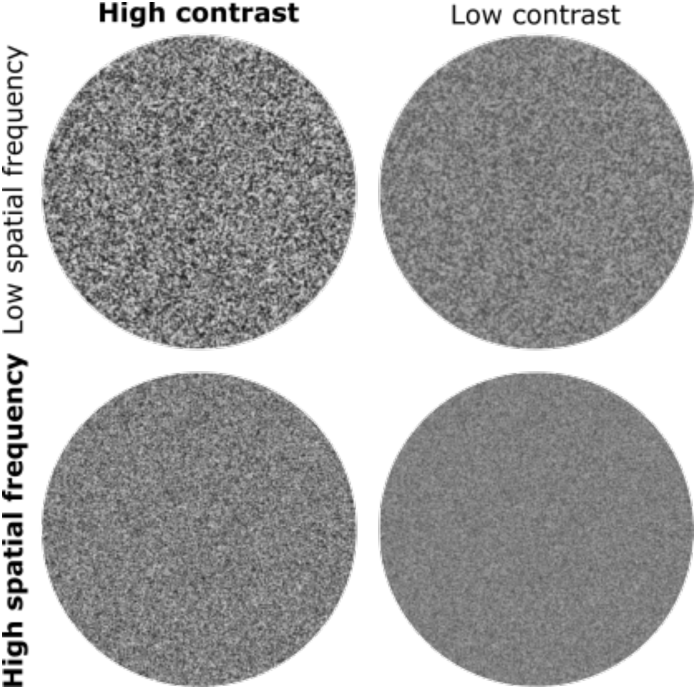
Some of the sand patterns conditions piloted. The select condition is shown in bold.

#### Auditory Steady-State Response

With respect to resonant properties in the auditory cortex, several studies have focused on gamma activity as inhibitory interneurons are thought to play a key role in the sensory processing differences in autism and related conditions (Edgar et al., 2016). These steady-state auditory evoked responses have also been found to vary with age (Arutiunian et al., 2023, 2025; De Stefano et al., 2019; Edgar et al., 2016; Wilson et al., 2007), making them valuable measures to study the development of auditory processing systems as they relate to autism.

In our protocol, we used a 500 Hz tone modulated in amplitude by either a 6 Hz or 40 Hz sine wave for 1s with a variable ISI (700-1300 ms range). The conditions (i.e., 6 Hz vs 40 Hz) were pseudo-randomized, with 20 repetitions of each condition. As with the Tone Oddball task, we originally had an abstract video playing in the background. However, we replaced these videos with the Disney short film *Luna* after we found it increased participant compliance. Although these more structured videos are susceptible to generating additional cognitive activity, we do not expect this activity to interact with the steady-state rhythms, being the variable of interest. The gain obtained by improving subjects’ compliance was judged to outweigh the risks of introducing such potential activations. During piloting, we experimented with replacing tones with white noise bursts but found tones to provide the best tradeoff between the salience of the neural responses and participant comfort. Pilot participants found tones more comfortable than noise, although the white noise stimuli provided higher response contrasts. The EEG responses to the 40 Hz modulation were the most conspicuous. The addition of a 10 Hz tone modulation was also piloted. However, we dropped this third condition to reduce the number of required trials and the task duration, to avoid potential interference from dominant alpha-band EEG responses, and to preclude the overlapping of the fourth harmonic of the 10 Hz condition with the fundamental of the 40 Hz condition. The 6 Hz tone modulation was maintained as a comparison condition to the 40 Hz stimuli as it also allowed comparing steady-state responses across modalities (visual and auditory) when compared to responses for the same frequency in the Visual Steady-State task. Further, this condition contained harmonics that were not overlapping with the 40 Hz condition (i.e., the 6th and 7th harmonics are at 36 Hz and 42 Hz and contain little power since power tends to decrease with increasing harmonic orders).

#### Naturalistic Social Preference

Development of the oculomotor system is a fundamental process that underlies higher-order attention and cognitive skills. Recent work using the CRISP (timer-**C**ontrolled **R**andom-walk with **I**nhibition for **S**accade **P**lanning) model suggests that fixation durations reflect online perceptual and cognitive ability in infants and are impacted by individual differences in the oculomotor system development (Saez de Urabain et al., 2017). Factors related to large between-individual variation in fixation durations are largely unexplored and may be relevant for understanding learning and oculomotor control in autism, particularly for social and non-social fixation preferences.

To address this gap, this task examines fixation durations and eye gaze patterns while viewing freely dynamic scenes. It relies on videos presented in naturalistic (normal) or semi-naturalistic (same as naturalistic but scrambled) conditions. We used 10 videos (5 in each condition) of 3 women performing baby-friendly actions (e.g., dancing with balloons) simultaneously or at different times. Each video lasts 21-25 seconds.

#### Pupillary Light Reflexes

The pupillary light reflex (PLR) is a reflexive physiological response that has been shown to differentiate autistic children and adults from neurotypical controls (Dinalankara et al., 2017; Fan et al., 2009; Nyström et al., 2018). In addition, retinal constriction has been found to correlate with atypical sensory processing in autistic children, suggesting that these low-level stimulus responses may be an important contributor to high-order sensory and perceptual processing deficits (Nyström et al., 2018). The PLR can also be used to study sympathetic and parasympathetic activations (Hall & Chilcott, 2018), providing an excellent approach to studying autonomic dysregulation in autism (Arora et al., 2021).

For this task, we used one block with 12 trials alternating a central fixation on a black background (0.9 lux; 6 seconds plus a 1600-2400 ms jitter) and a white background (190 lux; 75 ms). ET is used to determine that the participant is looking at the fixation point during the one-second window preceding the stimulus. The stimulus is not emitted if the participant is not looking at the fixation point. During piloting, we reduced the number of trials from 40 to 12 because this task was too uncomfortable for the participants and average responses were very clear even with this lower number of trials. This choice was also supported by other studies obtaining significant experimental results using even lower numbers of trials (Daluwatte et al., 2013; Fan et al., 2009; Soker-Elimaliah et al., 2022).

#### Visual Search

In this task, a target object is embedded in similar items as a standalone (e.g., a single red apple among blue apples) or in conjunction with distractor features (e.g., a full red apple among full blue apples and red apple slices). Several studies suggest that autism is associated with enhanced visual attention skills in some contexts, particularly visual search. This enhanced performance in autism is found across developmental stages and symptom levels (Kaldy et al., 2016). Developmental evidence suggests that enhanced perceptual performance during visual search is linked to the emergence of autism, such that visual search capacity in the first year of life predicts greater autistic symptoms between 1-2 years of age (Gliga et al., 2015). For this task, we used one block of 30 trials, arranged in a factorial design with three different numbers of distractors (5, 9, 13) and three mismatch conditions (color, size, conjunction). Corresponding numbers of repetitions are listed in Table 2. Trials were recycled if the reaction time for the gaze to the target was less than 100 ms (e.g., the participant was already looking where the target appeared) or larger than 2000 ms (the participant was not attentive). Preliminary analysis of the results from this task shows that the distance from the fixation point to the target has a large impact on the visual search time. As this distance has not been balanced across conditions, data analysis for this task requires controlling this distance. Future projects interested in reusing this task description are recommended to alter the protocol to balance this distance across conditions through pseudo-randomization.

**Table 2.** Number of trials per experimental condition. The conjunction condition combined mismatches in both colors and shapes.

| Distractors | Repetitions |  |  |
| --- | --- | --- | --- |
|  | Color mismatch | Shape mismatch | Conjunction |
| 5 | 2 | 2 | 6 |
| 9 | 2 | 2 | 6 |
| 13 | 2 | 2 | 6 |

### 2.4 Sample size

Although the recruitment is still ongoing, to date, the number of participants who completed the Q1K EEG/ET battery varied between 344 and 587 (Table 3), with the number of probands varying between 89 and 200, depending on tasks. Data from one relative is available for more than half of the proband participants, with data available for two relatives in about 10% of the cases. These numbers constitute a snapshot taken when this manuscript was written (July 8^th^, 2026) and are used for the analyses presented in the *Results* section. They are still, however, expected to increase with recruitment.

**Table 3.**
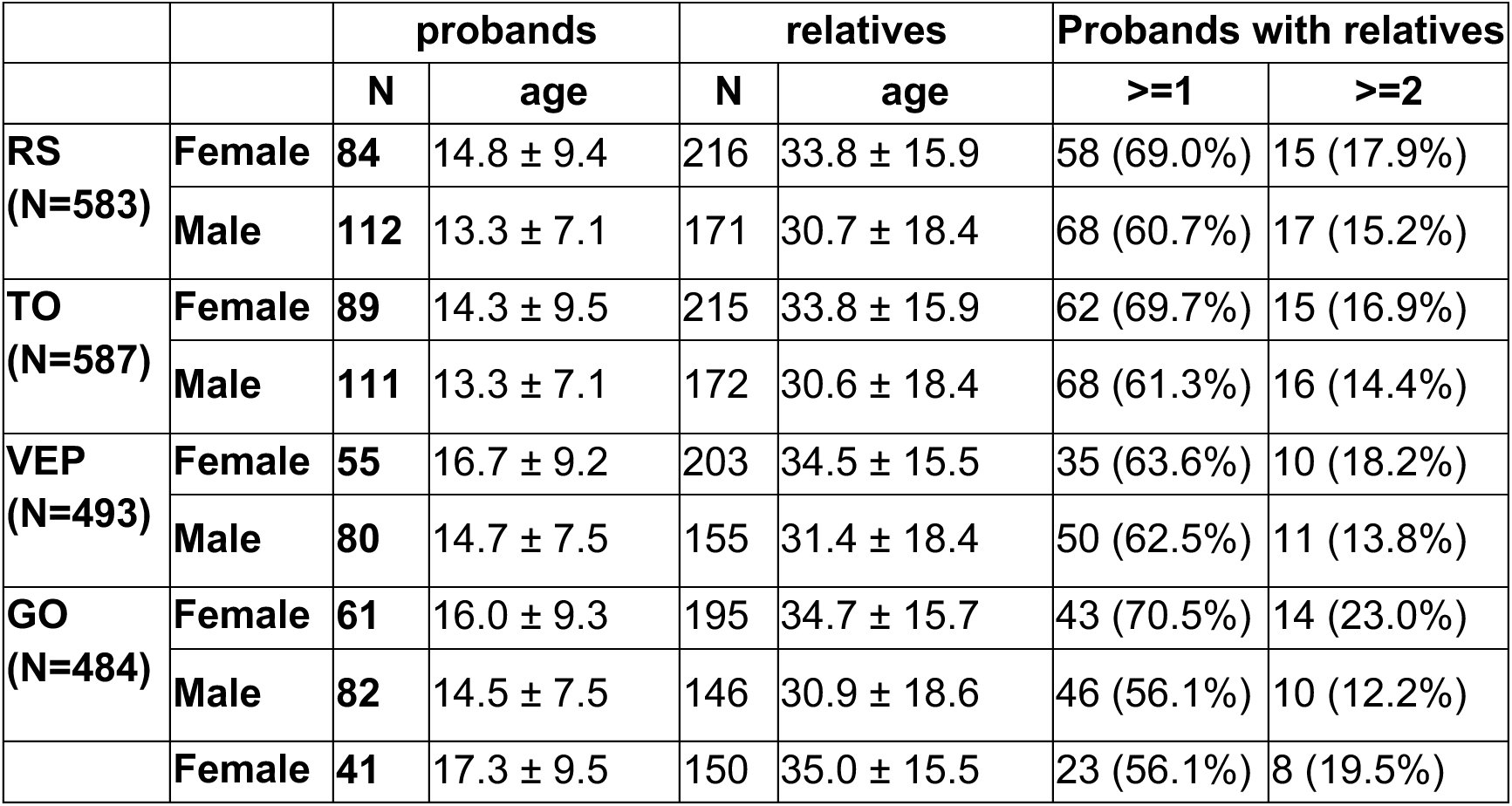
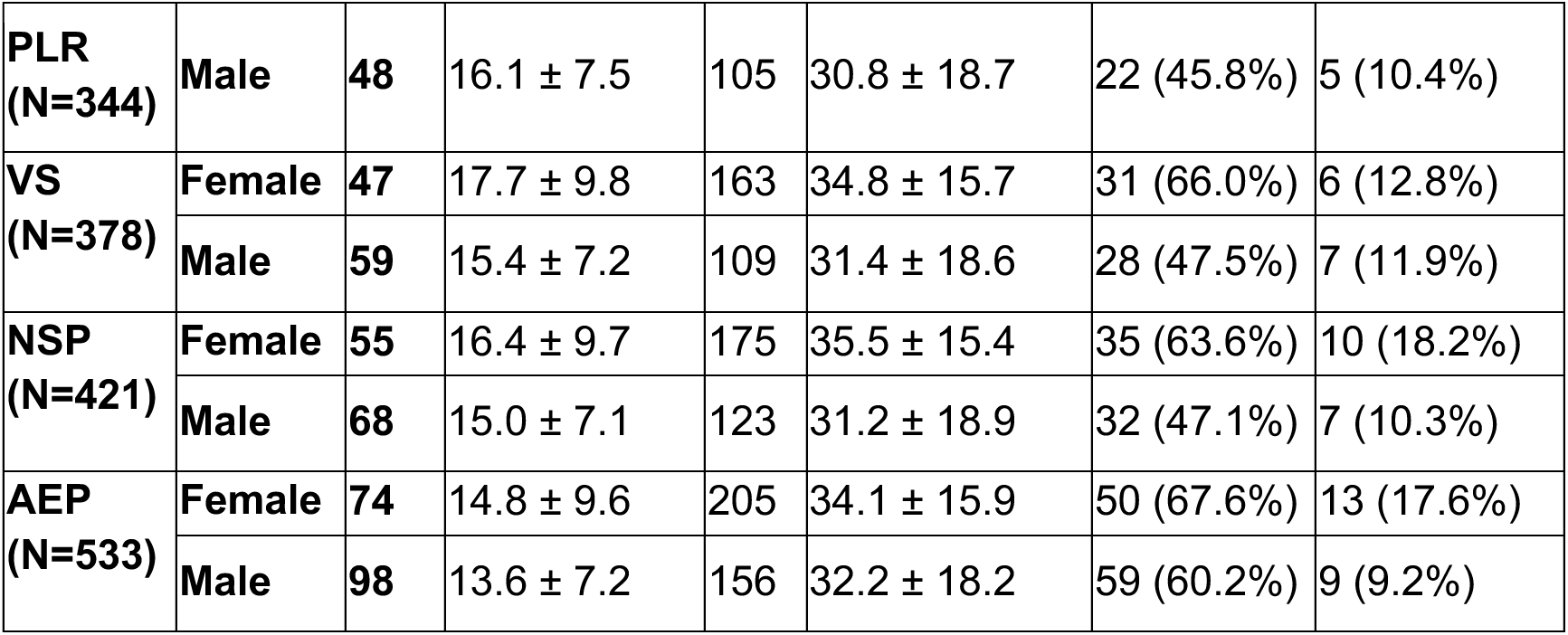
Number of participants per task and their split in probands and relatives.

### 2.5 System Integration

For this project, we designed an integrated EEG/ET acquisition system (Figure 2). In this architecture, the experimental protocol is run by the *Display PC*, a Windows machine running Experiment Builder (SR Research). The *Host PC* runs the acquisition software for the Eyelink 1000 Plus eye tracker system (SR Research). The iMac computer runs Net Station 5 for signal acquisition from a Net Amps 400 EEG amplifier (Magstim EGI). Both the EEG and the ET systems communicate with the acquisition software through the Ethernet protocol and are interoperable. To ensure high reliability and confidence in the timing of events reported by these systems, we further connected a Cedrus StimTracker to detect the timing of changes in luminosity on the screen and sounds emitted by the speakers, and record these events in Net Station and in the Eyelink software as *DIN*^3^ events. Changes in light intensity on the screen are detected by two photodiodes placed in a corner of the screen. The display for the different tasks has been designed with two small squares in this corner of the screen, which change from white to black with relevant task events, for example, with every onset-offset of the stimuli in the Visual Steady-State task. These events allow precise offline synchronization of the data stream from the EEG and ET systems with software like MNE-Python (Gramfort et al., 2013). We created a website containing more technical information to support external teams wanting to duplicate this setup: https://q1k-neuroimaging.readthedocs.io/en/latest/. Tutorials also demonstrate how to use MNE-Python to synchronize and integrate the EEG and the ET signals into a single EDF (European Data Format) file. However, for easier reuse, collected data are also shared in this integrated EEG/ET state, as described in the next section. This integration allows full flexibility to correlate eye-movement-related behaviors and neural activity, as demonstrated in the *Results* section. It also provides rich information to correct eye-movement-related artifacts (O’Reilly & Huberty, 2025, 2026) and disambiguate neural correlates of eye movement from higher-order cognitive processes (e.g., planning).

**Figure 2.**
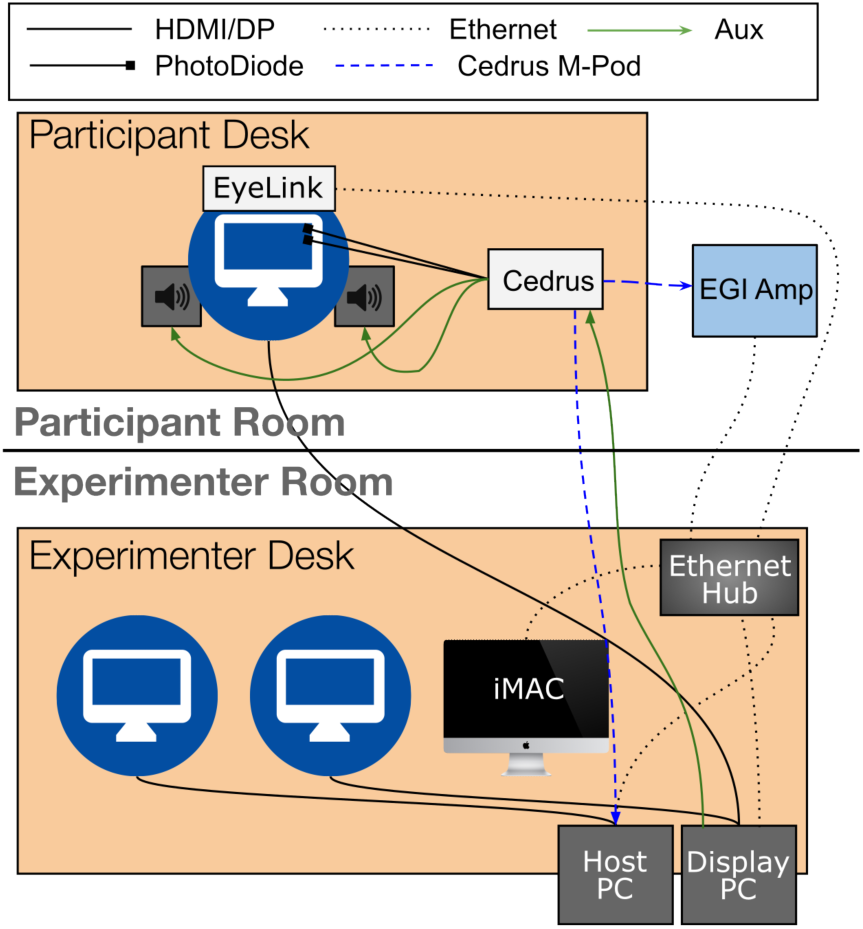
Diagram summarizing the integration of the ET and the EEG systems, including the addition of the stimulus tracker (Cedrus) to record events when changes in luminosity are detected on the screen or when sounds are emitted by the speakers.

### 2.6 Data Preparation and Management

#### Neuroinformatics

We implemented a pipeline for data management to ensure a smooth transition from acquisition sites to Open Access sharing (Figure 3). EEG/ET data are currently being collected at three sites: the Centre Hospitalier Universitaire de Sainte-Justine, Université de Montréal (HSJ), the Montreal Neurological Institute, McGill University (MNI), and the Centre Intégré Universitaire de Santé et de Services Sociaux du Nord-de-l’Île-de-Montréal (NIM). These sites also collect demographic data (collected and archived through RedCap) and other modalities (e.g., genetics, behavioral scores; collection and archival not displayed). Neuroimaging (7T MRI) is collected at the MNI site for all participants. We are currently setting up the software infrastructure to archive all these data types together on the Clinical, Biological, Imaging, and Genetic Repository (C-BIG) (Das et al., 2022), at the MNI. This repository includes a LORIS interface (Das et al., 2012) for managing neuroimaging and other clinical data. LORIS has recently been extended through the EEGNet project to support EEG data (manuscript in preparation). We previously worked to extend MNE-Python to allow the integration of ET signals within EEG EDF files as additional biosignals.

**Figure 3.**
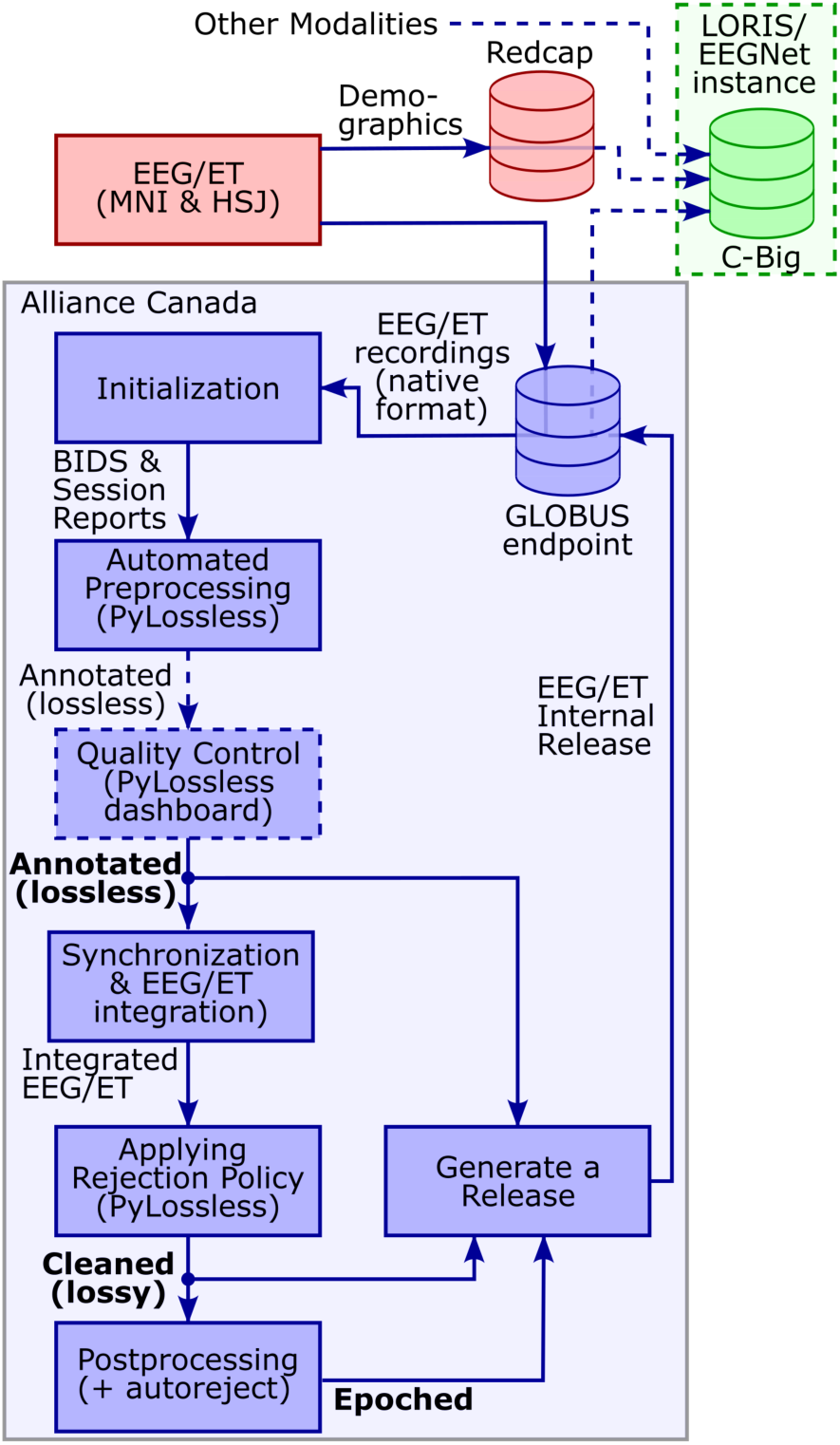
Data processing and generation of data releases for open sharing. Dashed paths are planned (not fully implemented yet). Bold fonts indicate shared data states (BIDS derivatives). Data are moved between infrastructures using a GLOBUS (Foster, 2011) endpoint.

#### Available data states

Before reaching C-BIG, EEG/ET data are augmented with annotations and metadata to maximize their usefulness to the scientific community. These manipulations are currently performed on Narval — one of the servers of the Digital Research Alliance of Canada — and lead to the preparation of an EEG/ET Internal Release. The generation of the internal release closely follows the philosophy of the PyLossless pipeline (Huberty et al., 2026) aiming at enriching the data with annotations without losing any information due to lossy transformations (e.g., filtering). All shared data are provided as Brain Imaging Data Structure (BIDS) datasets (Gorgolewski et al., 2016). We aim to share three data states: *annotated*, *cleaned*, and *epoched* (i.e., bold font data states in Figure 3). The annotated dataset is lossless, i.e., biosignals have not been modified from the recording version^4^. However, we augmented the dataset with annotations (noisy channels, time windows, and independent components) and cleaning procedures (e.g., rules for filtering, channel interpolation, and rereferencing). In this state, the dataset contains all the information required to generate a cleaned version of the recordings, but is still in a lossless format. This allows analysts to edit the cleaning process depending on their needs (e.g., change filtering parameters, avoid rejecting EOG artifacts, etc.) or to reprocess the datasets to benefit from new developments in preprocessing techniques. This state is ideal for people developing methods (e.g., preprocessing pipelines), integrating Q1K data with a dataset preprocessed with a different pipeline, or preferring to use different preprocessing pipelines. However, most analysts prefer leaving the cleaning of external datasets to the researchers who collected them and who have an intimate knowledge of the dataset’s properties. These analysts can use the annotated dataset and generate a clean (lossy) version by applying the available PyLossless Rejection Policy. However, to reduce this hassle, we also provide this cleaned, ready-for-analysis version of the dataset. Finally, in some cases, researchers prefer the data to be further preprocessed into a cleaned and segmented format. For example, this is often the case for applied machine learning researchers who may not be experts in EEG but want to develop classifiers or regression models to solve specific problems related to EEG analysis. For these people, the epoched format is provided.

#### Data release process

These three formats are integrated into internal releases whenever significant changes are made to the data. In line with the Q1K Open Science orientation, we aim to release data as early as possible. However, preprocessing and data cleaning is always improvable. Further, modification of a shared dataset may be required to keep up with changes in software environment and norms, for example, to integrate changes in a new version of the BIDS standard. To address these opposite requirements of early distribution and ongoing improvements, we adopted the concept of data release for this project. This approach allows both replicability and cyclic improvements. The internal EEG/ET release is pushed to C-BIG to be integrated with the other modalities (i.e., MRI, behavioral data, demographics, and clinical data). Whenever relevant, integrated multimodal releases can be archived on open access data repositories (e.g., the Canadian Open Neuroscience Platform (CONP; Poline et al., 2023)) to help with provenance tracking by distinguishing the latest modified version of the data available on the live CBIG repository from official “frozen” and versioned releases. C-BIG allows managing data through LORIS/EEGNet, which provides a web-based graphical user interface to perform additional quality control, data tagging using Hierarchical Event Descriptors (Bigdely-Shamlo et al., 2016), crowd-sourcing of annotations, and other similar data augmentation processes. The implementation of this releasing process is currently in progress.

## 3. Results

To demonstrate the quality of the collected data and exemplify the range of analyses made possible by our state-of-the-art EEG/ET integration, we show results for three tasks (Visual Steady-State Task, Pupillary Light Reflex, and Gap Overlap), not focusing on autism-related observations but fundamental properties of corresponding tasks.

### 3.1 Timing precision

To validate the reliability of the acquisition system timing and to assess potential site differences, we computed the inter-trial coherence (ITC) between epochs of the Visual Steady-State task (Figure 4). ITC is particularly sensitive to experimental timing because small timing offsets can break the coherence of the response across trials. Results in Figure 4 were computed with N= 266 participants for the HSJ site and N=30 for the MNI. We did not used the data from the sites with the smaller sample size (NIM; N=7) for this comparative analysis. The right-most column shows the between-site differences. Cluster-based permutation shows what is probably a false positive for the 6 Hz condition after the experimental stimulation. More significantly, the lower row shows how well the protocol captures the 6 Hz and 15 Hz entrainment and their multiple harmonics. The first three harmonics are all strongly statistically significant after an activation for the first few hundred milliseconds. This initial response is visible across frequencies, but with larger amplitude in low frequencies. It is visible for individual conditions but not for their contrast, suggesting that this initial activation is an evoked-related response to the stimulus onset, unrelated to the stimulation frequency. These results illustrate the quality of the data that will be available to the community to study the neural activation in people with autism and their family members.

**Figure 4.**
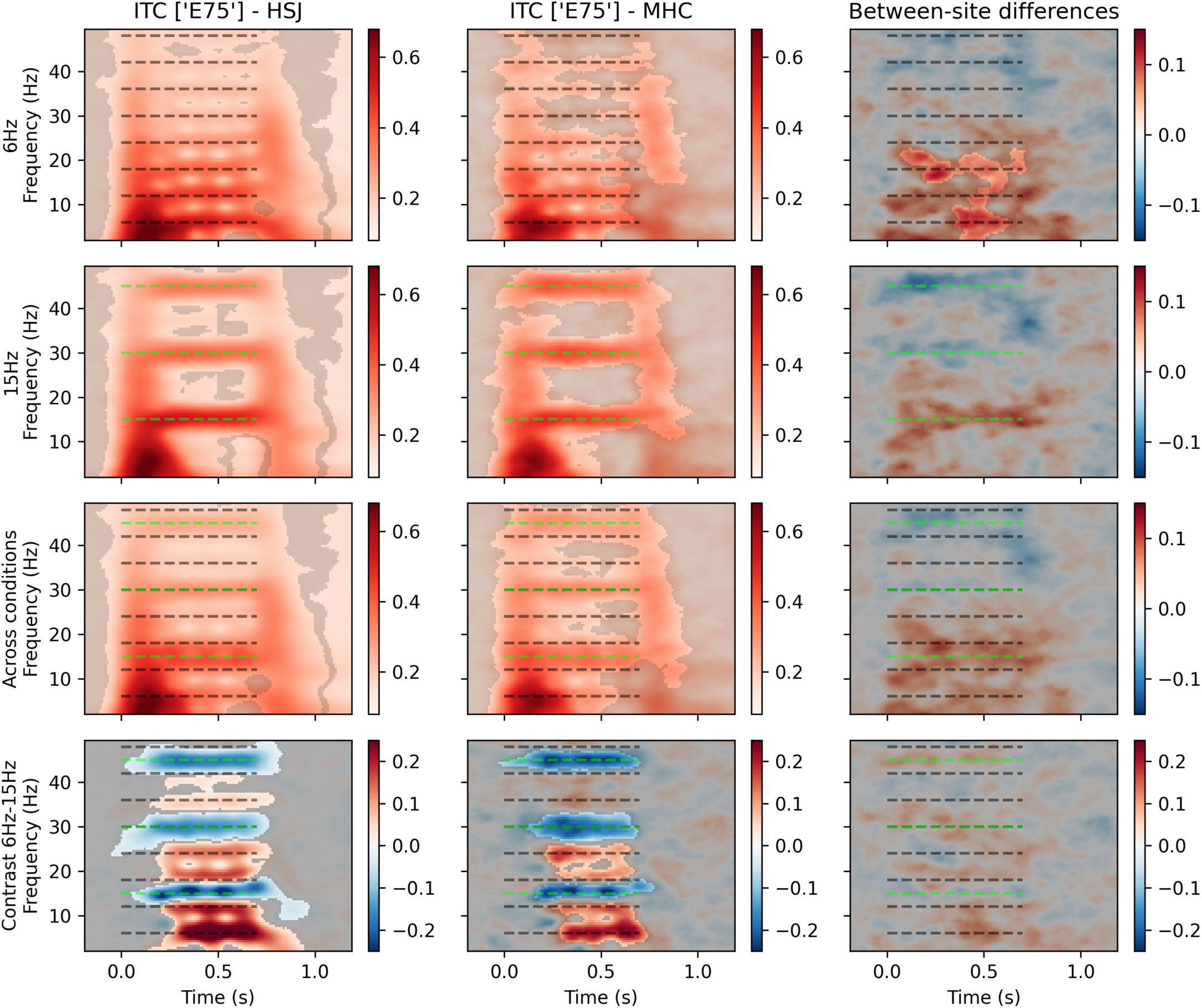
Inter-trial coherence (ITC) for the Visual Steady-State Task. The first two columns show the results for our two sites, while the third column shows the between-site difference. A shaded mask indicates statistical significance. The first two rows illustrate the ITC for the 6 Hz and the 15 Hz conditions, the third row gives the ITC averaged across conditions, while the last row shows the contrast for the ITC in the 6 Hz and 15 Hz conditions. Sample sizes: HSJ=266; MNI=30.

The strong effects observable in these preliminary data are essential for future studies interested in individual differences. Such investigations require very low experimental variability and high reliability to reach signal-to-noise ratios high enough to infer statistical effects within individuals.

### 3.2. EEG/ET integration and synchronization

We used the PLR task to demonstrate the potential provided by the integration of ET data (pupil size and gaze coordinates) directly in EEG EDF files. Although the PLR task is typically seen as an eye-tracking task, integrating EEG allows us to study event-related potentials associated with this protocol (Figure 5). Individual trials for a typical participant are illustrated in Figure 5.a,b. Individual average responses and overall average responses across all participants are illustrated in Figure 5.c-e. To identify central processing associated with the PLR task, we correlated the pupil response with the ERP for every channel and reported them as a topographical map (Figure 5.f). Cluster-based permutation was used to identify areas that are correlated above chance. This figure was prepared using the data from N=153 participants for the HSJ site, N=99 for the MNI site, and N=6 for NIM site. Given the association between the pupil response and the autonomic response, this dataset will allow researchers to study the relationship between autonomic and central activity and its potential modulation by autism, as demonstrated in Figure 5.

**Figure 5.**
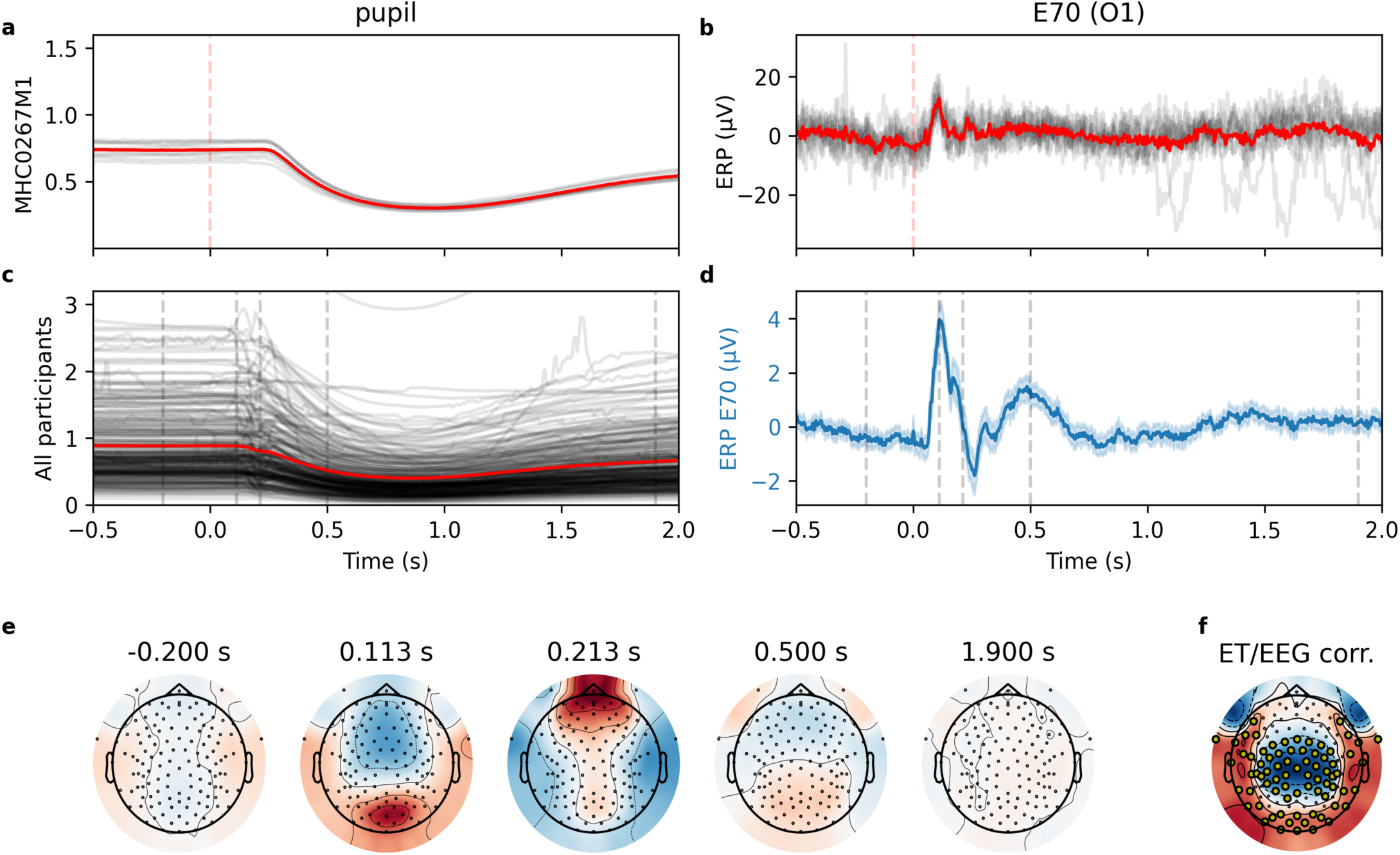
**a-b**) Example of pupillary and EEG responses for a typical participant (pale gray: individual trials; red: average responses) during the Pupillary Light Reflex task. c) Individual average pupil responses (gray) and overall average across participants (red). d) Average ERP for channel E70 (O1) with the 95% confidence interval (shaded). e) Topographical maps for times shown by dashed gray lines in panel d. f) Average correlation between the average pupil response and the ERP. Larger dots highlight significant clusters (p<0.01) using cluster permutation with one-sample t-tests. Sample sizes: HSJ=153; MNI=99; NIM=6.

Besides allowing the direct study of the relationship between ET and EEG signals through correlations or more sophisticated techniques (e.g., phase synchronization analysis), the EEG/ET integration provides eye-movement-related time markers for event-related EEG analyses. We used the Gap Overlap task to demonstrate this capability by computing ERPs for the three conditions (overlap, baseline, gap) when synchronizing with the stimulus onset and the gaze onset (Figure 6). Interestingly, the amplitude of the ERP obtained when synchronizing on the gaze onset is about twice the amplitude of those synchronized on the stimulus onset, showing that internally generated events captured through eye-tracking (i.e., gaze onset) may provide a more reliable way to synchronize brain responses for ERP analyses than the usual paradigm relying on the time of stimulus presentation. Such synchronization is available only in setups like the one we designed for Q1K.

**Figure 6.**
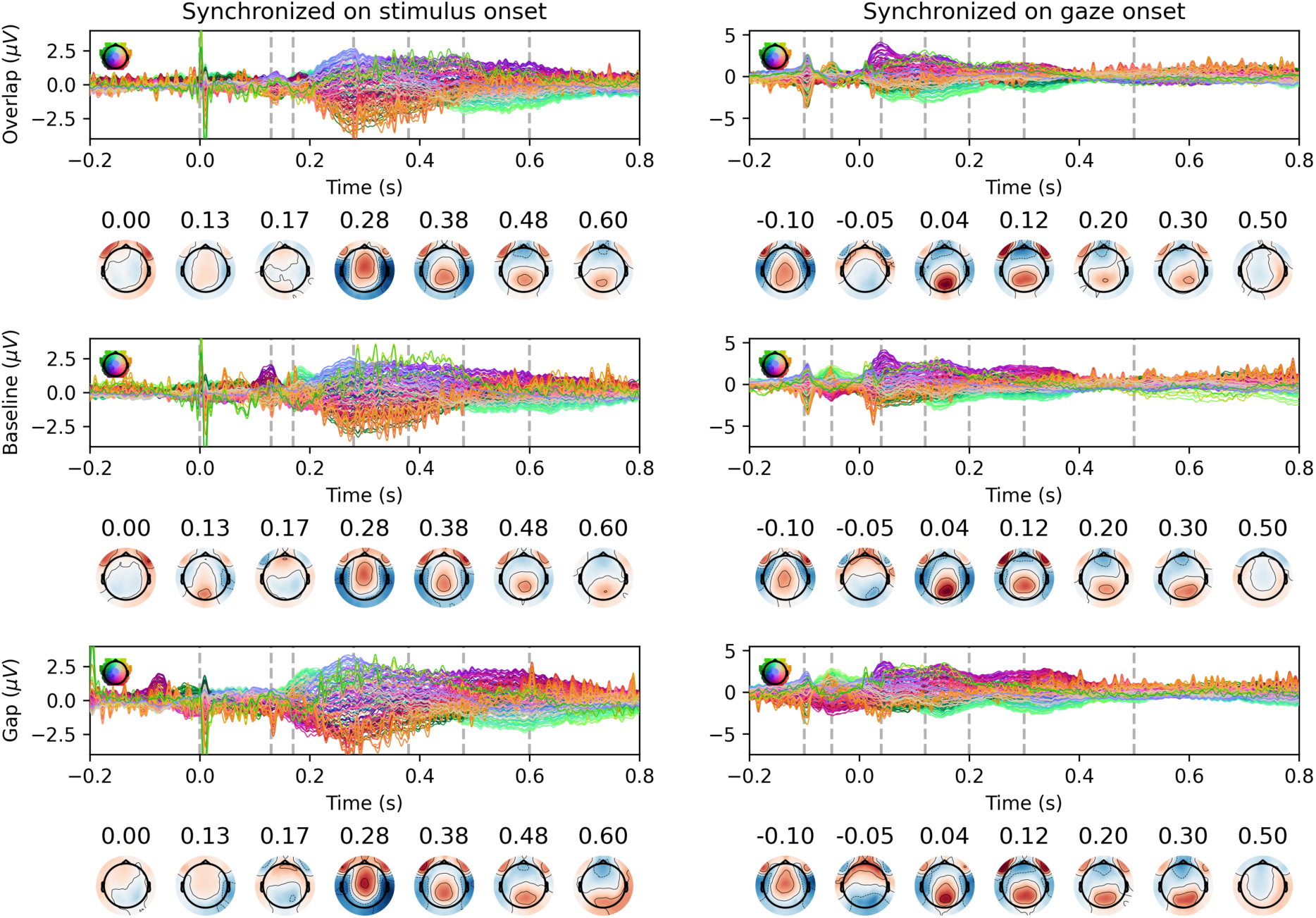
Gaze to peripheral stimuli showing the ERP upper panels and topography of EEG activity at different times (topomap strips) as displayed by dashed vertical lines on the ERP plots. Six such panels are displayed in a 3 x 2 grid, with the three rows being for the overlap (top), baseline (middle), and gap (bottom) conditions, and two columns depending on the synchronization on the stimulus onset (left) or the gaze onset (right). Sample sizes: HSJ=213; MNI=118, NIM=6.

Figures 4-6 illustrate only some of the novel types of analyses made possible with this new autism-related dataset. For example, for the Naturalistic Social Preference task, the availability of ET data allows for various types of novel ways to look at EEG response in autism in more naturalistic contexts than usually possible.

## 4. Discussion

The Q1K project has been designed within the Open Science framework. We are currently preparing the infrastructure to release this dataset openly. This dataset already includes more than 500 participants, and new participants are still being added. Importantly, although this paper focused on the EEG/ET data, Q1K adopted a deep, multimodal phenotyping strategy. Genetic, behavioral, and clinical data have also been collected on these participants. Neuroimaging with a 7T MRI scanner is also ongoing. Altogether, this diverse set of modalities provides a very rich dataset to investigate autism phenotypes.

Here, we are making available to the community the experimental protocol we designed (as EventBuilder files) (O’Reilly et al., 2026). The software infrastructure we implemented to convert the source files into BIDS format, preprocess them using the PyLossless pipeline (Huberty et al., 2026), synchronize and integrate EEG and ET, and postprocess them in epoched data has been packaged into an open-source Python library for easier reuse (https://github.com/lina-usc/q1k) and will be described in further detail in a manuscript in preparation.

Autism is a spectrum condition combining under the same umbrella people with different symptom profiles and etiologies. This large sample heterogeneity hinders the investigation of its mechanisms, as significant effects tend to wash away in diverse samples if such effects are not present in all strata. However, non-diverse samples may reveal effects that fail to be reproduced by other samples representative of different strata of the spectrum. Large datasets allowing rich phenotypical characterization are essential to develop our capacity to stratify autism samples by subtypes, to ensure that we cover the whole spectrum, and to adjust for differences in strata representation when comparing results across studies.

Previous studies have characterized different aspects of Q1K, but these findings have often remained fragmented across individual features, datasets, or analytical perspectives. By providing a thorough and integrated characterization, we aim for Q1K to support a more comprehensive understanding of the system as a whole, rather than a set of disconnected descriptions of its individual components. Further, by aiming to cover the complete spectrum and include participants with a wide Intellectual Quotient range, we hope to contribute open-access data allowing the characterization of autism in segments of the spectrum traditionally underrepresented in research.

## Data Availability Statement

The process for sharing openly Q1K data is currently being implemented. Q1K aims to make these data widely available for research in autism.

## Code Availability

The code for running this battery in Experiment Builder is freely available on Zenodo at https://zenodo.org/records/21268718. The code for data release infrastructure discussed in section 2.6 is available at https://github.com/lina-usc/q1k and is the objective of a manuscript in preparation.

## Authors Contributions

C.O’R. wrote the original manuscript draft, prepared the visualizations, developed software, contributed to methodology, performed formal analysis, and conducted the investigation. J.D. contributed to software development, methodology, and data curation. G.B.-G. contributed to software development and methodology. S.H., S.R., and J.B. contributed to software development. A.H.P.L., D.S., I.K., C.-O.M., and S.v.N. contributed to methodology. N.H.-L. and F.S. contributed to project administration. T.S., S.L., and M.E. contributed to conceptualization. B.F.d’A., C.E., M.Ca., C.L.T., J.S., M.Co., P.A., R.J., S.J., G.R., S.L., and M.E. acquired funding. S.L. and M.E. supervised the project. J.D., G.B.-G., S.v.N., B.F.d’A., J.S., R.J., S.J., S.L., and M.E. reviewed and edited the manuscript. All authors reviewed the manuscript.

## Competing Interests

The authors report no conflict of interests related to this work.

## Funding

The data and/or materials and/or registry resources used for this research were made available by the Quebec 1000 families (Q1K) project and through funds from the Fondation Marcelle et Jean Coutu and the Fonds de Recherche du Québec – secteur santé. This research was also enabled in part by support provided by Calcul Quebec (calculquebec.ca) and the Digital Research Alliance of Canada (alliancecan.ca).

## Acknowledgement

The authors wish to thank all families who contributed to the Q1K project. We acknowledge that linguistic preferences when talking about autism varies. We aligned the language of this document with the standard proposed in (Autism Alliance of Canada, 2025).

## Supplementary Information

**Supplementary Table 1.** ACAR clinic equipment.

| Hardware | Software | Specifications |
| --- | --- | --- |
| Electrical Geodesics EEG Net<br>Amps 400 amplifier | N/A | Sampling rate: 1000 Hz |
| Electrical Geodesics Hydrocel<br>129 channel Net | N/A | 129 channel (128 channels + Cz reference channel) net with Eye channels (125-128) removed. |
| Cedrus Stimtracker | N/A | Photocell + 3mm audiojack,<br><u>Photocell 1:</u><br><u>Photocell 2:</u><br><u>Audio L:</u><br><u>Audio R:</u> sensitivity must be set at 3 |
| iMac | Net Station 5 acquisition<br>software (Mac OSX) | 27 inch display<br>OSX 10.13.4<br>Intel Core 2 Duo Processor (2 Ghz)<br>16 GB RAM |
| SR Research Eyelink 1000<br>Plus Eyetracker ("Host PC") | Eyelink 1000 Plus Host<br>software v5.15 | Monitor Dimensions:<br>29 cm x 18 cm x 9 cm<br>Sampling rate: 1000 Hz<br>Resolution: 0.05° RMS<br>Saccade resolution: 0.25°<br>Trackable range: 32° x 25°<br>Lens: 25mm |
| Ultra Performance PC<br>("Display PC") | Experiment Builder 2.2.299<br>DataViewer<br>Windows 10 64-Bit |  |
| ASUS VG248QE, 24"<br>Participant Display |  | screen size: 530 mm x 300 mm<br>screen resolution: 1920 x 1080 |
| Polhemus FASTRAK | Brainstorm Digitization<br>Software | Standard FASTRAK system with :<br>● 1 X 4 inch source<br>● 1 X 8 inch stylus<br>● 3 X RX2 sensors |
| USB-RS-232 adapter | N/A | USB to RS232 Adapter with PL2303 Chipset<br>USB 2.0 Male to RS232 Male Gold Plated DB9<br>Serial Converter for Windows 10, 8.1, 8, 7, Vista, XP, 2000, Linux and Mac OS X 10.6 and Above<br>Brand: CableCreation |

**Supplementary Table 2.** Protocol workflow events. Typically, not required for analysis.

| Description | EGI Event |
| --- | --- |
| begin task | VBeg |
| Net Station misc. event | TSYN |
| display start menu | dstr |
| display break menu | dbrk |
| display end screen | dend |

**Supplementary Table 3.** Events for the Resting-State task.

| Description | EGI Event | DIN |
| --- | --- | --- |
| begin task | VBeg |  |
| Display Video | vs0[1,2,...,6] | DIN3 |
| Net Station misc. event | TSYN |  |
| display start menu | dstr |  |
| display break menu | dbrk |  |
| display end screen | dend |  |

**Supplementary Table 4.** Events for the Tone Oddball task.

| Event Description | EGI Event |  | DIN |  |
| --- | --- | --- | --- | --- |
| Condition | Standard | Deviant | EGI | Eyelink |
| Onset of tone | mmns | mmnt | DIN4 | 4 |

**Supplementary Table 5.**
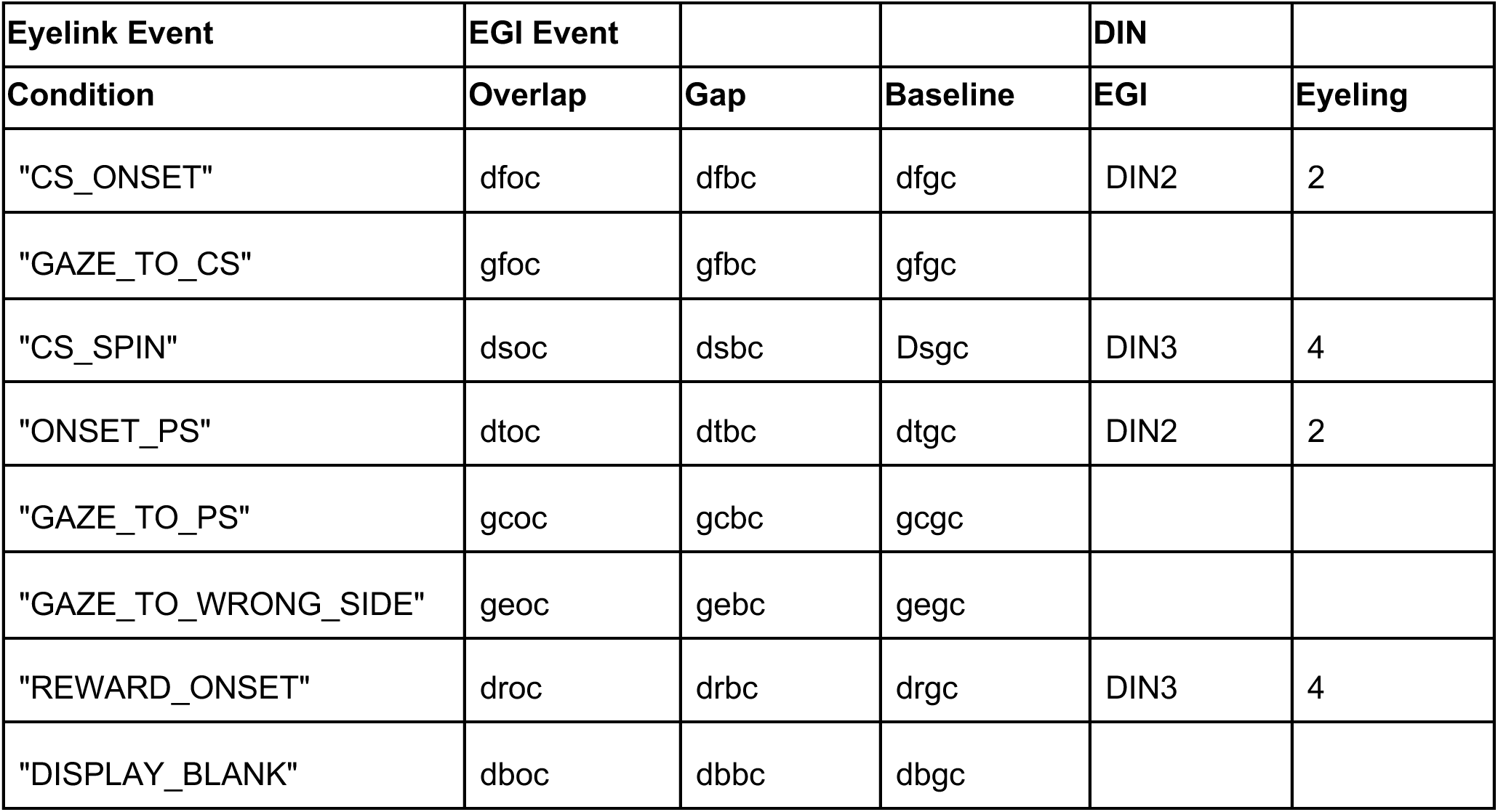
Events for the Gap Overlap task.

**Supplementary Table 6.** Events for the Visual Steady-State task. This task also has four fixation animation onset events: fvsr, fvsb, fvct, and fvcr.

| Event Description | EGI Event |  | DIN |
| --- | --- | --- | --- |
| Condition | 6 Hz | 15 Hz |  |
| Fixation animation onset | fvsr | fvsb |  |
| Onset of 1-second video | sv06 | sv15 | DIN2 |

**Supplementary Table 7.** Events for the Auditory Steady-State task.

| Event Description | EGI Event |  | DIN |
| --- | --- | --- | --- |
| Condition | 6 Hz | 40 Hz |  |
| Onset of 1-second tone | ae06 | ae40 | DIN7 |

**Supplementary Table 8.**
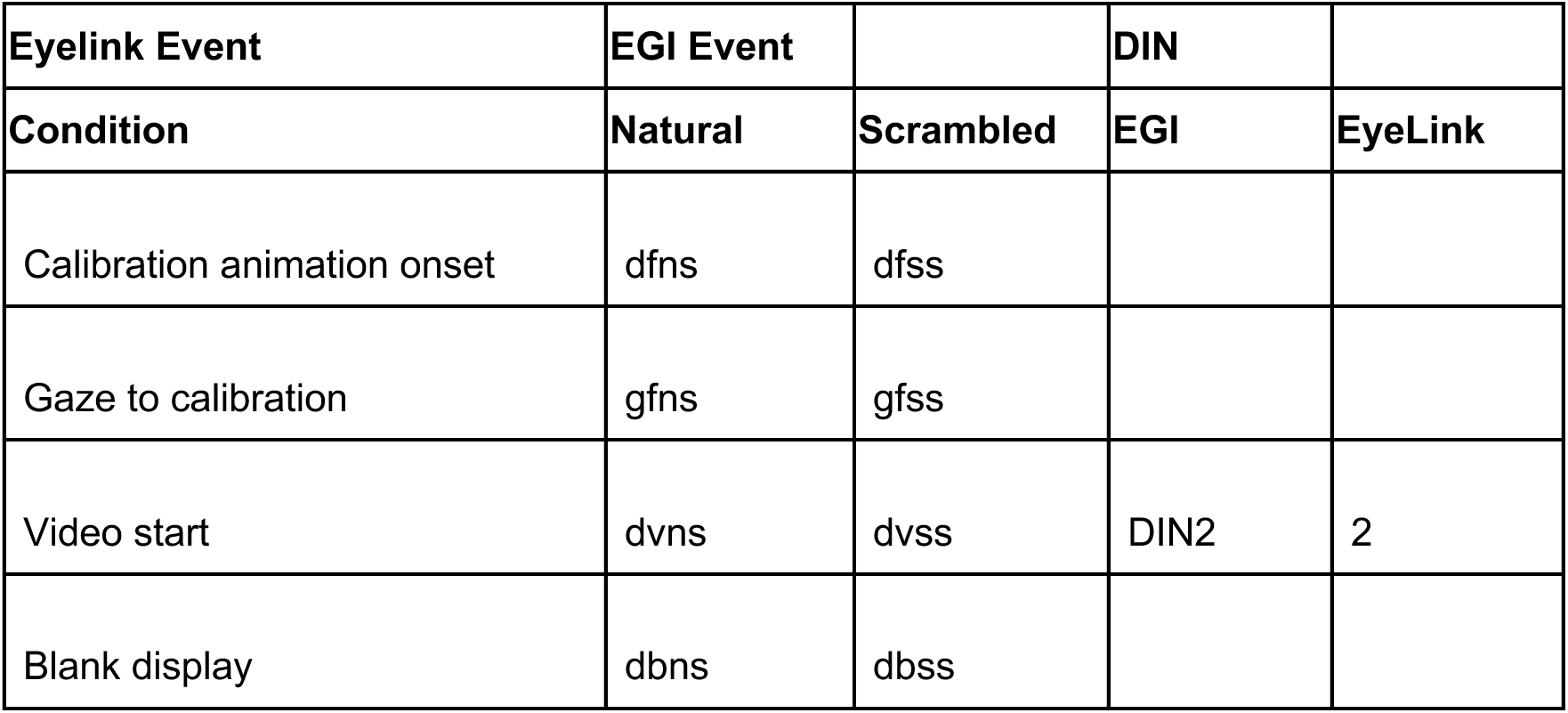
Events for the Naturalistic Social Preference task.

| **Eyelink Event** | **EGI Event** |  | **DIN** |  |
| --- | --- | --- | --- | --- |
| **Condition** | **Natural** | **Scrambled** | **EGI** | **EyeLink** |
| Calibration animation onset | dfns | dfss |  |  |
| Gaze to calibration | gfns | gfss |  |  |
| Video start | dvns | dvss | DIN2 | 2 |
| Blank display | dbns | dbss |  |  |

**Supplementary Table 9.**
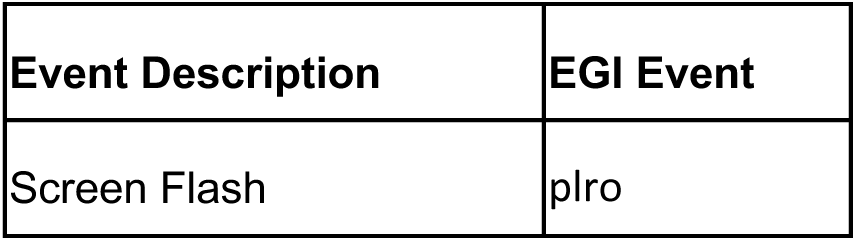
Events for the Pupillary Light Reflexes task.

**Supplementary Table 10.** Events for the Visual Search task.

| **Event Marker** | **Red Apple Slices** |  |  | **Blue Whole Apples** |  |  | **Conjunction** |  |  | **DIN Event EGI** | **DIN Event EYELINK** |
| --- | --- | --- | --- | --- | --- | --- | --- | --- | --- | --- | --- |
|  | **5** | **9** | **13** | **5** | **9** | **13** | **5** | **9** | **13** |  |  |
| Apple fly in | da5s | da9s | datc | da5a | da9a | datc | da5c | da9c | datc | DIN3 | 4 |
| Gaze to attention apple | ga5s | ga9s | gatc | ga5a | ga9a | gatc | ga5a | ga9a | gatc |  |  |
| Display fixation | df5s | df9s | dftc | df5a | df9a | dftc | df5c | df9c | dftc | DIN2 | 2 |
| Display search | ds5s | ds9s | dstc | ds5a | ds9a | dstc | ds5c | ds9c | dstc | DIN3 | 4 |
| Gaze to target | gt5s | gt9s | gttc | gt5a | gt9a | gttc | gt5c | gt9c | gttc |  |  |
| Display reward | dr5s | dr9s | drtc | dr5a | dr9a | drtc | dr5c | dr9c | drtc | DIN2 | 2 |
| Display blank | db5s | db9s | dbtc | db5a | db9a | dbtc | db5c | db9c | dbtc |  |  |

## Footnotes

1 Including younger participants for the 7T MRI protocol would require the development of a pediatric coil due to minimal weight constraints in high fields. Efforts are underway to address this limitation, and the inclusion of younger participants may be possible in the future.

2 The obtention of extremely clear inter-trial coherency patterns is demonstrated in Figure 4, in the Results section.

3 DIN refers to *Deutsches Institut für Normung* which established a standard for connectors and gave its name to these connectors. When a change of voltage is detected on one of the pins of such a DIN connector, it logged as a DIN event.

4 More accurately, minimal modifications (e.g., low and high-pass filtering) are currently applied by PyLossless in the generation of its “lossless” state. This issue is planned to be resolved in future versions of PyLossless to ensure a truly lossless state.

## Notes

### Competing Interest Statement

The authors have declared no competing interest.

